# Droplet rather than Aerosol Mediated Dispersion is the Primary Mechanism of Bacterial transmission from Contaminated Hand Washing Sink Traps

**DOI:** 10.1101/392431

**Authors:** Shireen Kotay, Rodney M. Donlan, Christine Ganim, Katie Barry, Bryan E. Christensen, Amy J. Mathers

## Abstract

An alarming rise in hospital outbreaks implicating hand-washing sinks has led to widespread acknowledgement that sinks are a major reservoir of antibiotic resistant pathogens in patient-care areas. An earlier study using a GFP-expressing *Escherichia coli* (GFP-*E. coli*) as a model organism demonstrated dispersal from drain biofilm in contaminated sinks. The present study further characterizes the dispersal of microorganisms from contaminated sinks. Replicate hand-washing sinks were inoculated with GFP-*E. coli*, and dispersion was measured using qualitative (settle plates) and quantitative (air sampling) methods. Dispersal caused by faucet water was captured with settle plates and air sampling methods when bacteria were present on the drain. In contrast, no dispersal was captured without or in between faucet events amending earlier theory that bacteria aerosolize from P-trap and disperse. Numbers of dispersed GFP-*E. coli* diminished substantially within 30 minutes after faucet usage, suggesting that the organisms were associated with larger droplet-sized particles that are not suspended in the air for long periods.

**IMPORTANCE:** Among the possible environmental reservoirs in a patient care environment, sink drains are increasingly recognized as potential reservoir of multidrug resistant healthcare-associated pathogens to hospitalized patients. With increasing antimicrobial resistance limiting therapeutic options for patients, better understanding of how pathogens disseminate from sink drains is urgently needed. Once this knowledge gap has decreased, interventions can be engineered to decrease or eliminate transmission from hospital sink drains to patients. The current study further defines the mechanisms of transmission for bacteria colonizing sink drains.

## INTRODUCTION

Recent reports have implicated hand-washing sinks as a primary reservoir of antibiotic resistant pathogens within patient-care environments(1–27). Many of these reports have been published since 2016 highlighting the global recognition that biofilms located in and on sinks can have in disseminating clinically important drug resistant gram-negative bacteria(14–27). Retrospective and prospective surveillance investigations affirm that hospital sinks provide habitats for several opportunistic pathogens, raising serious concerns(8, 24, 26, 28). It is not the mere presence of these drug-resistant pathogens in the hospital wastewater that is of concern, but the ability of these organisms to colonize biofilms on the luminal surfaces of wastewater plumbing and thereby withstand routine cleaning practices. While several gammaproteobacteria detected from the sinks in hospitals have been linked to healthcare-associated infections, opportunistic pathogens like *Pseudomonas aeruginosa, Acinetobacter baumannii* and *Stenotrophomonas maltophilia* are typically known to be found in water environments(29–31). In contrast, emerging pathogens such as the carbapenemase-producing *Enterobacteriaceae*, many having fecal origin, may survive within the biofilm formed on sink surfaces and wastewater premise plumbing(32, 33). Often by acquiring mobile resistance elements through horizontal gene transfer, carbapenemase-producing *Enterobacteriaceae* (CPE) infections are especially threatening because they are more frequent causes of highly antibiotic resistant infections with reduced treatment options.

In outbreak investigations species and strain matching between patient and sink isolates is often attributed to sink source contamination, however direction (sink to patient versus patient to sink) and precise mode of transmission remains inconsistent and elusive(29, 30). Even with increased recognition of transmission, a knowledge gap exists with regards to the precise mechanism of transmission from sink reservoirs to the patient. Using a model of sink colonization with green fluorescent protein (GFP)-expressing *Escherichia coli* (GFP-*E.coli*), we recently demonstrated the source and the degree of dispersion from sink wastewater to the surrounding environment(34). Factors effecting the rate and extent of droplet-mediated dispersion were investigated, but particle size involved in dispersion was not measured in this study. Studies that claim aerosols as the primary dispersion mechanism from sinks are based on rudimentary findings (2, 23, 35, 36) or assumptions drawn based on these unsubstantiated findings(3, 6, 10, 13, 21, 37). Airborne particles originating from sinks can have varied sizes and compositions. The World Health Organization and Healthcare Infection Control Practices Advisory Committee (HICPAC) guidelines use a particle diameter of 5 μm to delineate between bioaerosol (≤5 μm) and droplet (>5 μm) transmission (38, 39).

Aerosol-mediated transmission and droplet-mediated transmission in the healthcare environment will require conceptually different infection control strategies. Clarity regarding aerosol versus droplet mediated dispersion in the context of sinks is critical. In the present study, we aim to further define several outstanding knowledge gaps: i) dispersion mechanism of bacteria, aerosol sized particles or droplets, from biofilms in handwashing sinks, ii) factors triggering dispersion from a colonized sink drain and iii) role of biologically active aerosols spontaneously dispersing from drain or P-trap without a triggering event. GFP-*E. coli* as the surrogate organism for Enterobacteriaceae was used in this model study to investigate these questions.

## MATERIALS AND METHODS

### Sink Gallery Operation and Automation

A dedicated sink gallery at the University of Virginia and described in an earlier study(34) was used in the present study. Sinks were operated in automated mode using a microprocessor. The microprocessor activated the faucets for 30 s each hour via an inline solenoid valve in the hot water supply line (faucet event) and also turned on a peristaltic pump (Masterflex Pump #HV-77120-42, Cole-Parmer, Vernon Hills, IL) to dispense 1ml soap (Kleenex Foam Skin Cleaner, Kimberly-Clark Worldwide Inc., Roswell, GA). A steel tube discharging the soap was held in a clamp attached to the faucet and positioned directly below the discharge of the faucet water. Water flow rate from the faucet was 8 L/min. In one experiment, mannequin hands (Dianne Practice Hand, D902, Fromm International, Mt. Prospect, IL) attached to a metal rack and positioned between the faucet head and sink drain were used (Figure 1). To facilitate access to the luminal surface of the drain line, sampling ports were drilled along the length of the tailpiece (between the P-trap and the drain), and the trap arm (between the P-trap and the common line). These holes were fitted with size 00 silicone stoppers (Cole-Parmer, Vernon Hills, IL). Temperature and total and free chlorine residual concentrations of the faucet water i) first catch and ii) after 2 min flushing of faucets were measured at regular intervals using the DPD method (Hach Model, Hach, Loveland, CO).

**1.**
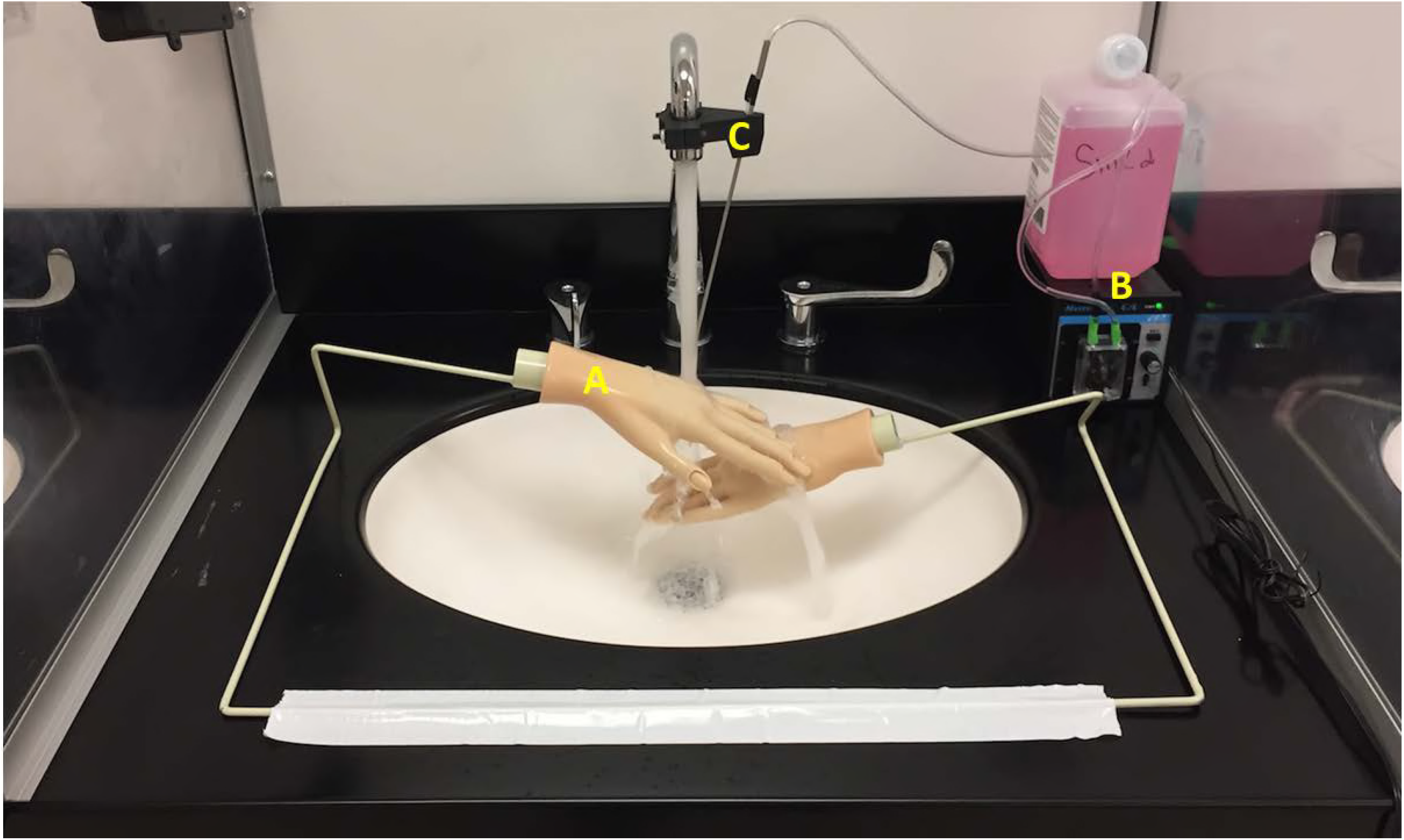
Test sink, showing mannequin hands positioned directly below faucet (A), and a peristaltic pump on the left to deliver hand-soap (B), via steel tube attached to the faucet (C).

### Inoculation, Growth and Establishment of GFP-*E.coli* in Sink P-traps

A single isolated colony of GFP-*E.coli* (ATCC^®^ 25922GFP^™^) grown from −80°C stock was inoculated in 5 ml Tryptic Soy Broth (TSB) containing 100μg/ml ampicillin (ATCC^®^ Medium 2855). The method of inoculation varied for each experiment. For P-traps a 10ml mid-log phase culture of GFP-*E.coli* (10^9^ CFU/ml) was added into the P-trap water (~150ml) through the lower-most sampling port on the tailpiece using a 60ml syringe attached to silicone tubing (Cole-Parmer, Vernon Hills, IL). The inoculum was mixed with the P-trap water by repeated withdrawal and injection of the inoculum, with precautions taken to avoid unintentional inoculation of drain (strainer) or sink bowl (bowl). For the drain inoculation, a 10ml mid-log phase culture of GFP-*E.coli* (10^9^ CFU/ml) was evenly applied on the surface of the drain using a sterile pipet. For establishment of GFP-*E.coli* P-trap biofilm, a 10 ml mid-log phase culture of GFP-*E.coli* (10^9^ CFU/ml) was added into an unused P-trap and following inoculation, 25ml TSB and 25ml (x2) 0.85% saline was added on a daily basis through the drain for 7 days to facilitate biofilm growth on the luminal surface of the drain line without additional water every hour. Seven days later, P-trap water and swab samples from the inner surface of the drain, tailpiece, and the P-trap were plated on Tryptic soy agar containing 100μg/ml ampicillin (TSA). TSA plates were incubated overnight at 37°C and colony-forming units (CFUs) fluorescing under UV light were enumerated. All preparatory culturing of GFP-*E.coli* took place in a separate room from the sink gallery to avoid unintentional contamination.

### Sampling and Enumeration of *GFP-E.coli*

To monitor the growth of GFP-*E.coli* within the plumbing, sterile cotton swabs (Covidien^™^, Mansfield, MA) presoaked in 0.85% sterile saline were inserted through sampling ports and biofilm samples were collected by turning the swab in a circular motion on the inner surface (~20 cm^2^). Sample swabs were pulse-vortexed in 3ml saline and serial dilutions were plated on TSA. The entire surface area of the drain, faucet aerator, and sink bowl surfaces were sampled using environmental sponge wipes (3M Sponge-stick with neutralizing buffer, 3M, St. Paul, MN) using overlapping and multidirectional motions. The sponge-wipes were expressed in 90 ml of Phosphate Buffered Saline containing Tween 80 (0.02%) (PBST) using a Stomacher (400 Circulator (Seward Ltd., UK). The eluate was concentrated by centrifugation and plate counts were performed on TSA and R2A agar (Becton Dickinson and Company, Franklin Lakes, NJ). TSA plates were incubated at 35 °C for 48hr and fluorescent CFUs were enumerated; R2A plates were incubated at 25°C for 7d and CFUs were counted.

### Experimental Approach for Dispersion Studies

Each dispersion experiment comprised a 30 s faucet event repeated three times at 60 min intervals (Fig. 2). Each experiment also comprised a control (without faucet events) followed by a test (with faucet events). Three sinks were tested concurrently but staggered by a few minutes, to account for the variability in dispersion driven by faucet water flowrate, air flow dynamics in the room, contact angle and wastewater drainage / water backup rate.

**2.**
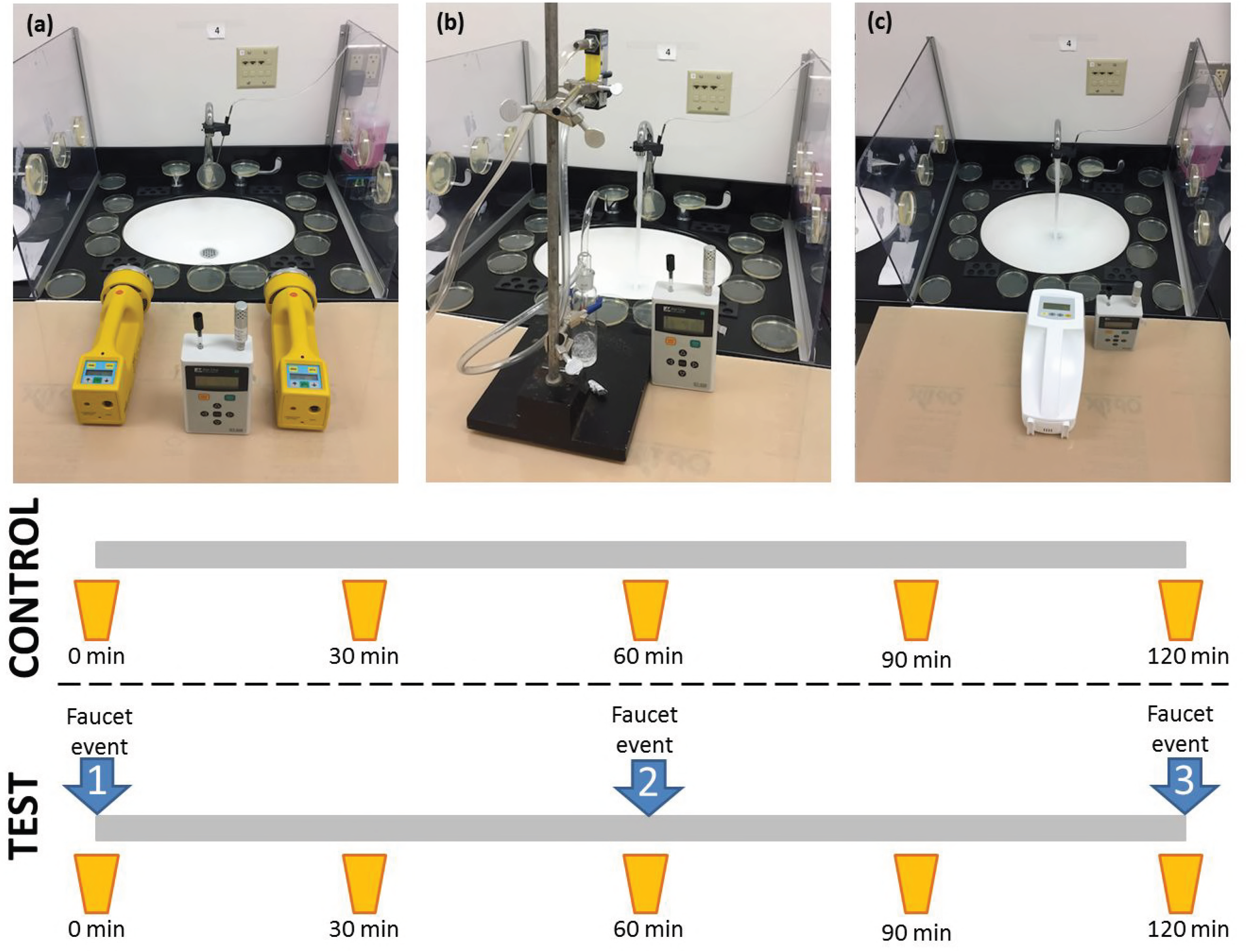
Experimental set-up used for different air sampling methods: a) Impaction, b) Impinger, and c) Gel Filtration. Air samples were collected at the initial faucet event (0 min) and every 30 minutes thereafter. Faucet events (faucet activation) occurred at 0, 60, and 120 minutes under test conditions. Faucets were not activated in control experiments.

### Sampling Droplet Dispersion

TSA settle plates were used to capture the droplet dispersion. Numbered TSA plates were laid out radially around the sink bowl. A fixed layout and number of settle plates around the sinks was used for each dispersal experiment (Fig. 3). The counter space of each sink was thoroughly disinfected with Caviwipes-1 (Metrex Research, LLC, Orange, CA) prior to each experiment. TSA plates were then positioned on the sink counter surrounding the sink bowl. Additional plates were attached to the faucets, plexiglass partitions, and faucet handles using adhesive tape. Plates were not placed in the sink bowl. TSA plates were also placed >3 m away from the sink as negative controls. Lids of the TSA plates were removed only for the duration of the dispersal experiment. Dispersion per defined area (CFU/cm^2^) for settle plates was determined by dividing the CFU counts in the TSA plate by the surface area of the plate.

**3.**
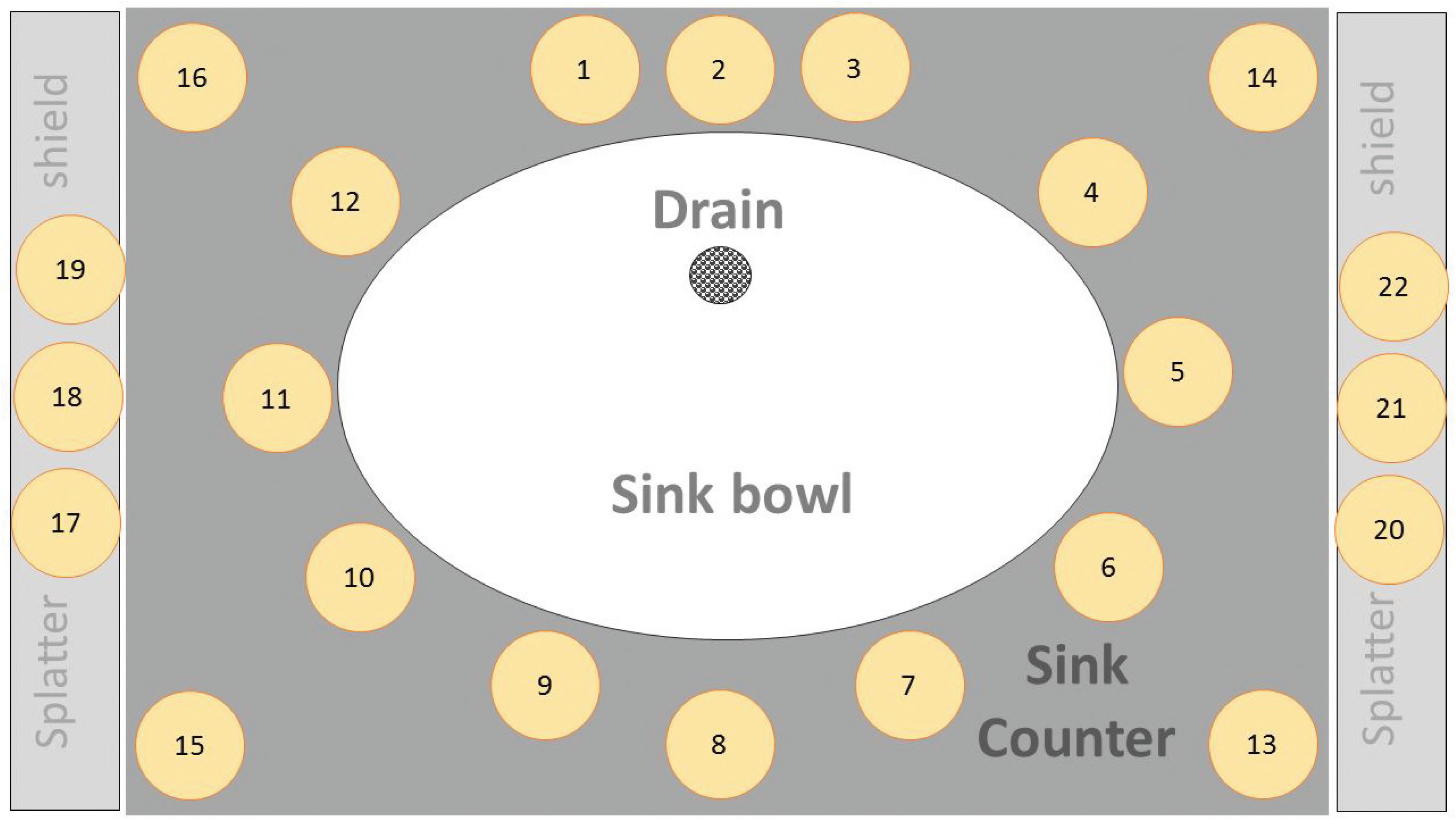
Graphical representation depicting the layout of the settle plates positioned around the sink used to capture droplet dispersion.

### Air Sampling and Particle Counts

Each dispersion experiment comprised the collection of air samples at five separate time points: time t=0 (first faucet event), t=30 minutes after the first faucet event, t=60 minutes (second faucet event), t=90 minutes, and t=120 minutes (third faucet event) (Fig. 2). Individual sink sampling was staggered by a few minutes to provide time for air sampler installation and sampling of each sink. A control experiment period, in which faucets were not activated for the entire 120 minutes was also performed for each sink. Three air sampling methods were tested: impaction, impingement, and filtration. For the impaction method, two SAS90 air samplers (Bioscience International, Rockville, MD) containing one TSA plate and one R2A plate each were positioned 12 inches from the sink bowl and set for a 300L sample (at 90L/min for 200 seconds) (Fig. 2a). TSA and R2A plates from each air sampling event were incubated as described earlier. A gel filtration device (MD8 Portable Air Sampler-Sartorius AG Goettingen, Germany) fitted with disposable gelatin filters (Sartorius AG Goettingen, Germany) was positioned 12 inches from the sink bowl and set for a 300L sample (at 100L/min for 180 seconds) (Fig. 2c). Gelatin filters were carefully overlaid on TSA plates, which were as already described. Liquid impingers (Ace Glass Inc. Vineland, NJ) were autoclaved and filled with 20ml sterile Phosphate Buffered Saline (PBS) prior to each experiment. Each was connected via a flowmeter (Cole-Parmer, Vernon Hills, IL) and vacuum pump (Cole-Parmer, Vernon Hills, IL). The impinger was positioned 12 inches from the sink bowl (set at 6L/min for 50min) to collect a 300L air sample (Fig 2b). In a biological safety cabinet, the liquid from the impinger was transferred to a sterile tube, vortexed, filtered through 0.22μm membrane filters (Pall Laboratories, Port Washington, NY), and 5 ml duplicate samples were plated on TSA and R2A plates. Fluorescent CFUs were enumerated after TSA plates were incubated at 35°C for 48 hours and counted. R2A plates were incubated for 7 days at 25°C and counted. Plates from air impaction samples and samples collected from liquid impingement were shipped via overnight courier to CDC laboratories for processing and counting. Gel filtration plates were processed and counted at University of Virginia. Paired with air sampling particles in size range 0.3, 0.5, 0.7, 1.0, 2.0 and 5.0 μm were measured using particle counter (GT-526, Met One Instruments, Inc. Grants Pass, OR) placed 12 inches from the sink bowl (Fig 2.). With a runtime of 660 seconds each, 3 successive runs of particle counter were performed, first run coinciding with t=0. Particle counter also recorded relative humidity and air temperature.

### Verification of GFP-*E. coli*

Fluorescent colonies on TSA-amp plates were counted under a long-wavelength UV light source. To verify that fluorescent colonies were GFP-*E.coli*, two fluorescent colonies from each sample were randomly selected and first screened on MacConkey II agar (BD, Franklin Lakes, NJ). The MacConkey II plates were incubated at 35°C for 24 hours, and lactose fermenters were isolated on Tripticase Soy Agar with 5% Sheep Blood (TSA II) (BD, Franklin Lakes, NJ) and incubated under the same conditions. Once colonies were isolated, they were identified using matrix assisted laser desorption ionization-time of flight mass spectrometry (MALDI-TOF MS) (Bruker, Billerica, MA), or using the Vitek 2 system (bioMérieux Durham, NC).

### GFP-*E.coli* Detection in Sink Plumbing and faucet water

Prior to each experiment, 500 mL of first-catch faucet water and a 500 mL sample collected after two minutes of flushing were collected in sterile bottles containing sodium thiosulfate (0.18 g/l) for dechlorination. Samples were plated on TSA and incubated at 35°C for 48 hours to test for GFP-*E.coli* in the faucet water supplied to each sink and to ensure cross contamination of GFP-*E.coli* had not occurred. First-catch and two-minute flush samples were diluted and plated on R2A, which were incubated at 25°C for 7 days and counted. After the experiment was completed, samples were collected and processed to detect GFP-*E.coli* in the sink plumbing. The P-trap water was collected, and processed by vortexing and filtration, and plated in duplicate on TSA and R2A media as already described. The sink P-trap and tailpiece were removed from the sink lab units, filled with faucet water, plugged, and shipped by overnight courier to CDC for analysis. Swab sampling from the tailpipe and P-trap, and sponge wipes were processed as already described to recover and quantify biofilm organisms. Samples were plated on TSA and R2A media, and counted, as described.

## RESULTS

Free and total chlorine concentrations in the faucet water were consistent across the experiments (Table 1). So were the water and air temperatures (Table 2). In comparison, the relative humidity recorded across the experiments varied, with highest recorded in case of P-trap inoculation experiments (Table 2).

**Table 1.** Total and Free Concentrations and temperature in the Faucet water. Mean and standard deviation of total and free chlorine residual and water temperature, measured at different time points and from water collected from faucets supplying different sinks over the course of this study.

|  | Initial (First Catch) | Final (After 2min) |
| --- | --- | --- |
| Total Chlorine | 0.77 ±0.26 mg/L | 0.63 ±0.07 mg/L |
| Free Chlorine | 0.62 ±0.25 mg/L | 0.54 ±0.13 mg/L |
| Water Temperature | 22.53 ±1.87°C | 36.69 ±2.17°C |

**Table 2.** Air temperature and relative humidity recorded across experiments.

|  | Air Temperature (°C) | Relative Humidity |
| --- | --- | --- |
| Drain Inoculation | 19.8 | 31.4 |
| P-trap Inoculation | 19.5 | 54.4 |
| Drain + P-trap Colonization | 20.0 | 44.7 |
| Control experiments | 19.7 | 46.5 |

### Dispersion immediately following P-trap inoculation

No GFP-*E. coli* dispersion was detected on settle plates immediately following inoculation of P-traps with 10^10^ CFU GFP-*E. coli* (Fig. 4a). No dispersion was detected using impaction, impingement, or filtration air sampling methods (Fig. 5). Further, no GFP-*E.coli* were recorded in the P-trap water, P-trap, tailpiece or sink bowl surface samples collected at the end of dispersion experiment.

**4.**
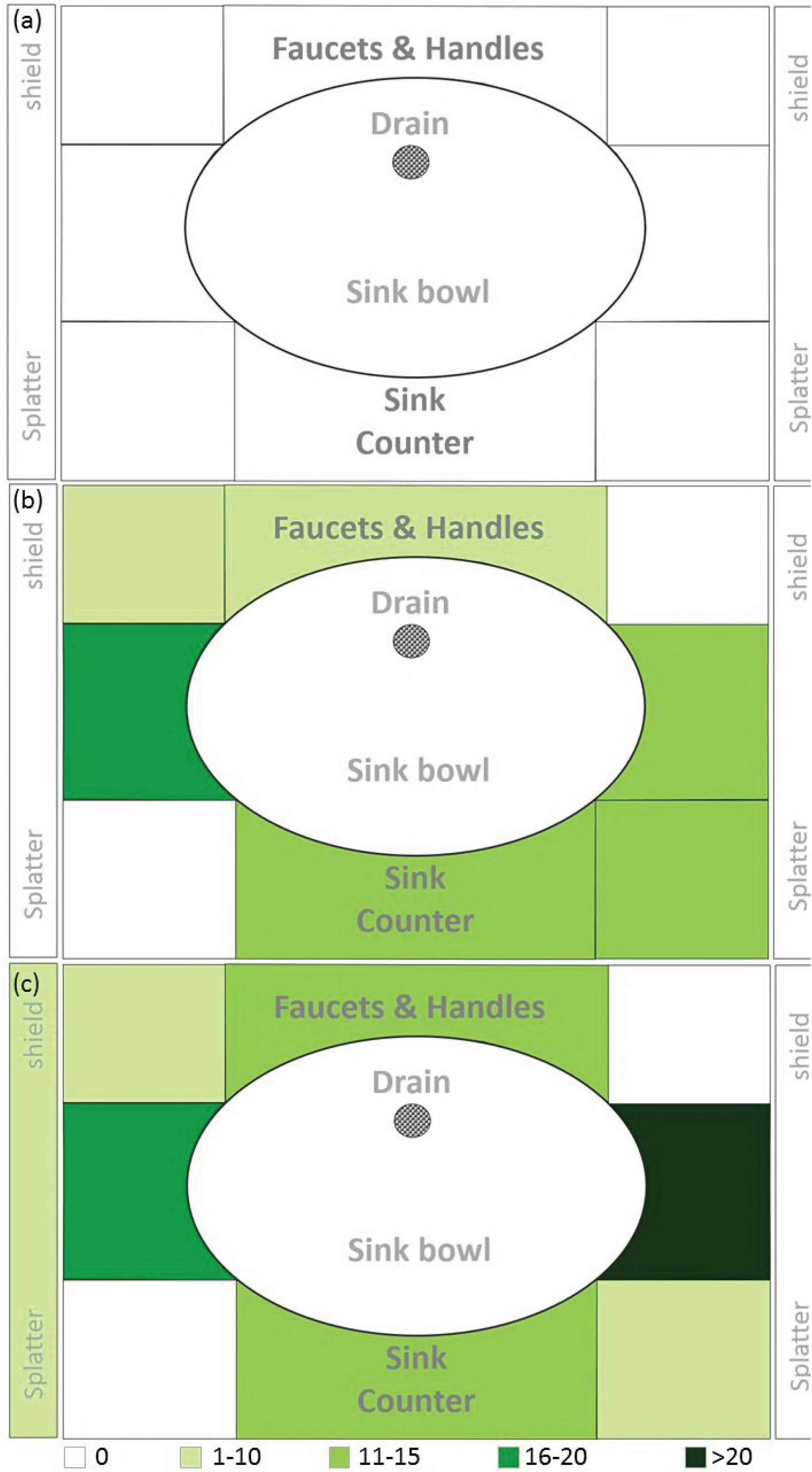
Heat map representation of GFP-*E.coli* dispersion captured on TSA settle plates following (a) P-trap, (b) sink drain inoculation and (c) drainline colonization.

**5.**
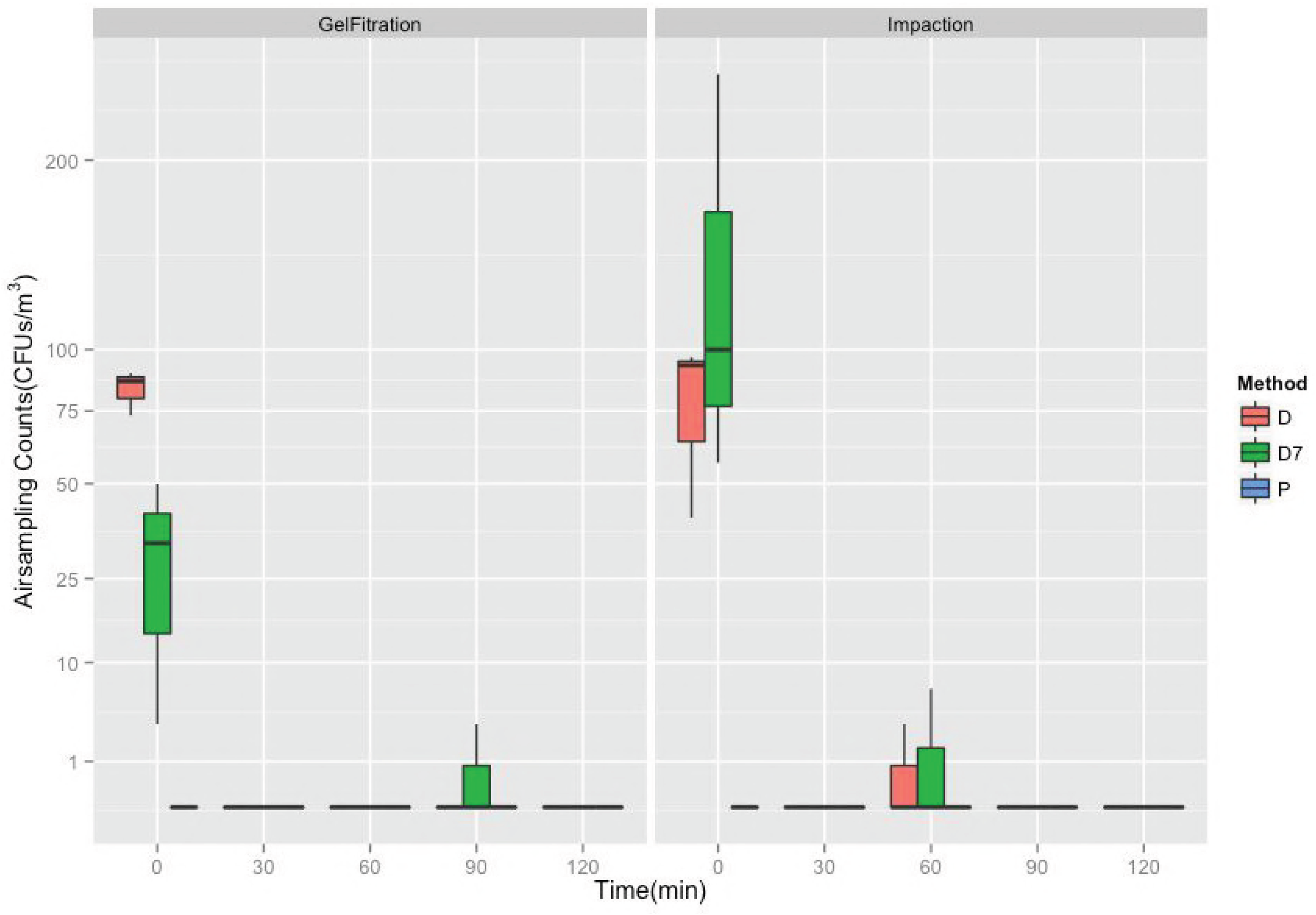
GFP-*E. coli* as measured by impaction [SAS90], and gel filtration [MD8] across D-Drain inoculation, D7-Drainline colonization and P-P-trap inoculation methods.

### Dispersion immediately following Drain inoculation

When sink drains were inoculated with 10^10^ CFU GFP-*E. coli*, dispersion was detected on settle plates and by impaction and filtration. Dispersion detected by settle plates across the three sinks ranged from 35-107 CFU/plate. GFP-*E. coli* levels were higher on the counter space surrounding the sink bowl compared to lower counts near the faucets (Fig 4b). Dispersion was not detected on side-splatter shields. With the exception of the detection of 1 CFU/m^3^ at t=60 min using the impaction, dispersion of GFP-*E. coli* was detected only at the first faucet event (t=0 min) using both impaction and filtration methods (Fig 5). Average dispersion captured at the first faucet event (t=0min) was 77 and 83 CFU/m^3^ using impaction and filtration methods respectively. GFP-*E. coli* was not detected at any time point using liquid impingement.

### Dispersion following growth for 7 days in an amended P-trap biofilm

Allowing colonization of GFP-*E. coli* in drainlines between strainer and P-trap with nutrient exposure over time, dispersion was detected on settle plates (Fig. 4c), with counts ranging from 49-107 CFU/plate. The counter space surrounding the sink bowl received the largest amount of droplet dispersion, followed by faucet, faucet handle surfaces and splatter shields. GFP-*E. coli* levels were highest at the first faucet event (t=0 min) and not detectable afterwards, with the exception of a 2 CFU/m^3^ count at 60 min using air impaction and 1 CFU/m^3^ count at 90 min filtration similar to what was observed for drain inoculation (Fig. 5). Dispersion captured at the first faucet event (t=0 min) was 138 and 29 CFU/m^3^ using impaction and filtration methods, respectively. GFP-*E.coli* was not detected using liquid impingement. GFP-*E.coli* was detected on the sink bowl, drain grate, tailpiece, P-trap, and P-trap water at the completion of this experiment (data not shown).

### Dispersion without faucet events (control experiments)

Without a faucet event, GFP-*E. coli* was not detected on settle plates, or by impaction, impingement, or filtration air sampling methods in any experiment. Without a faucet event fungal and non-fluorescent CFUs were occasionally recorded on settle plates, subsequently identified as *Staphylococcus sp*., non-hemolytic Streptococci, *Paenibacillus sp*., yeast and small gram positive rods.

### Total viable heterotrophic organisms in laboratory air

No GFP-*E. coli* was detected in the faucet water at any time during experiments. Air samples were collected by impaction on R2A medium with and without faucet events, in order to quantify total heterotrophic organisms in the air space in proximity to the sink during each experiment. Across the experiments the heterotrophic organisms in the air ranged from 4–578 and 2–69 CFU/m^3^ with (test) and without (control) faucet events, respectively (Fig. 6). Dispersion captured at the first faucet event (t=0min) when the faucets were turned on (test) was 1 log_10_ higher than the same recorded in case of control experiment (without faucet event). Heterotrophic organisms captured from the air steadily declined along the time points 0, 30, 60, 90 and 120 min with and without faucet events. Particle concentrations in the air with and without faucet events were found to be consistent during the day of the experiment. However, when compared across the experiments, no correlation could be established (Supplemental Figure S1).

**6.**
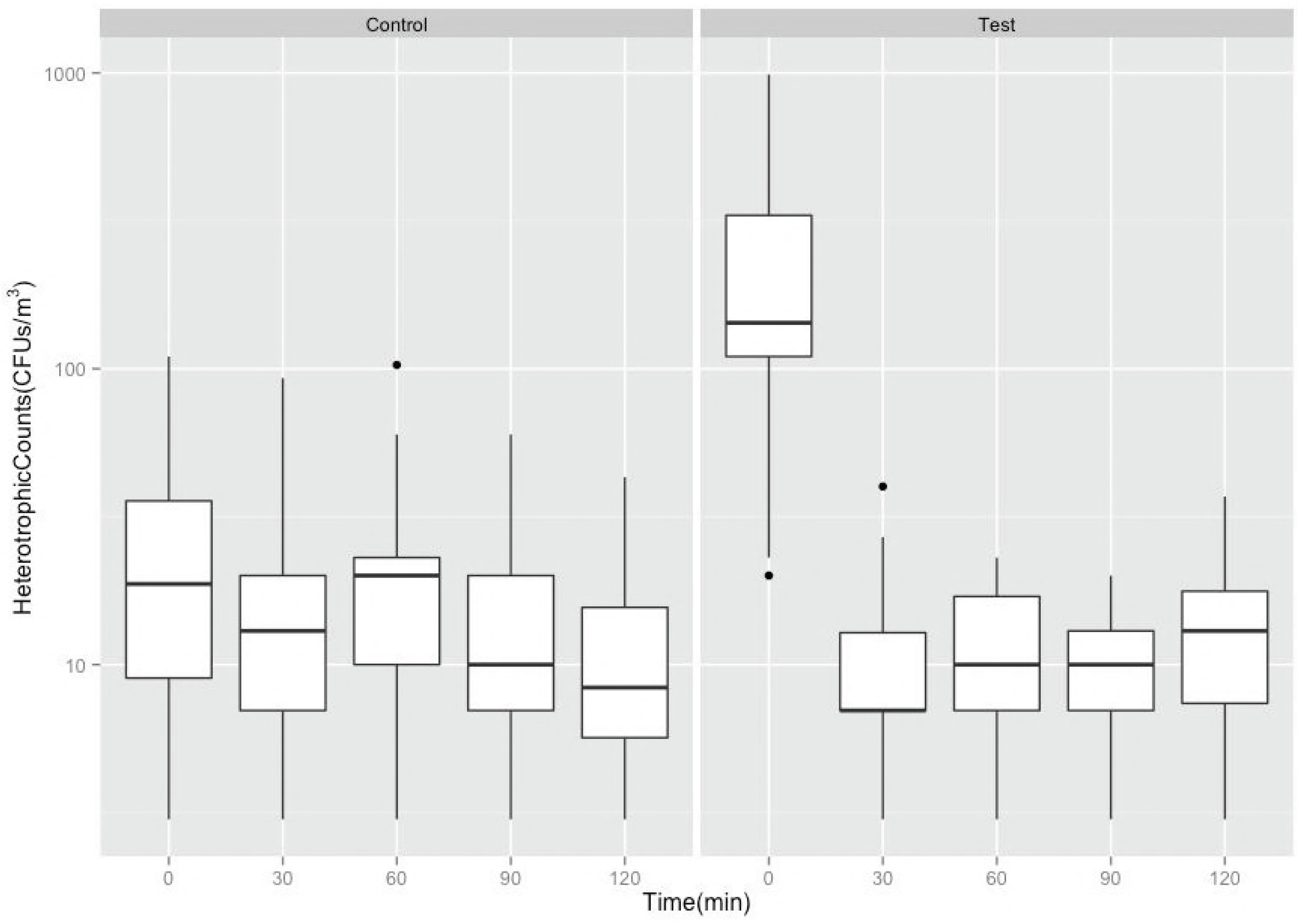
Heterotrophic Plate Counts as measured by impaction air sampling a) without faucet event (Control) and b) with faucet events (Test).

### Dispersion in the Presence of Mannequin Hands

When mannequin hands, which created a barrier for water flow onto the drain, were positioned under the faucet and the sink drain was inoculated with ~1E+10 CFUs GFP-*E.coli*, 10-fold lower dispersion was captured on settle plates. No dispersion in air sampling using impaction or filtration methods was observed. Dispersion pattern and load on settle plates varied considerably across the three sinks and experiments with the mannequin hands in place.

## DISCUSSION

The objective of the present study was to characterize the mechanism of bacterial dispersion from handwashing sinks, using a GFP plasmid-containing *E. coli* strain as a surrogate for multidrug resistant Enterobacteriaceae. Very few studies have investigated the dispersion from sinks using methods to sample aerosol-associated microorganisms(23, 36, 40, 41); however, several studies have drawn subjective interpretations about aerosol-mediated transmission from contaminated sinks(3, 6, 10, 13, 21, 37). Doring et al. used an impaction method to examine *P. aeruginosa* dispersion from the sink bowl surface during faucet use. *P. aeruginosa* was detected 15 cm from the sink drain when counts in the sink drains exceeded 10^5^ CFU/ml (41). Kramer et al. detected 439 CFU/m^3^ *P. aeruginosa* in the air sampled 10cm above the sink drain with 10^5^ CFU/ml bacteria in the ‘sink fluid’ (P-trap water)(40). In these studies, dispersion was not measured without faucet usage (control samples). De Geyter et al. used an MAS-100 air sampler to measure CRE dispersal from contaminated sinks with and without faucet usage. Several species in the family Enterobacteriaceae were detected during faucet usage but results for control samples (without faucet usage) were not provided(23). Fusch et al. in contrast sampled air around sinks with and without faucet event (control) and did not detect *P. aeruginosa* in air samples collected without running water(36). We had previously provided a quantitative assessment of dispersal as a function of faucet usage and reported GFP-*E. coli* could be dispersed up to 30 inches beyond the sink drain during faucet usage(34). However, dispersion was assessed using a gravity method, and bioaerosol production was not evaluated.

The air sampling methods chosen and tested in the present study were three of the most widely used methods previously reported(23, 35, 36, 40, 42, 43) and were selected to assess bioaerosol production during sink usage. In the present study, GFP-*E. coli* dispersion was detected during a faucet event but was not detected in the absence of faucet events using either settle plates or impaction and filtration air sampling methods. This finding corroborates previous studies (2, 21, 34, 36). It also implies that shear forces of the faucet water flowing onto the sink drain and/or bowl surfaces results in dispersion of bacteria. Detection of dispersed GFP-*E. coli* during a faucet event and non-detection at subsequent time points (after 30 minutes) suggested that dispersed cells were associated with larger heavier droplets that would quickly settle onto surfaces due to gravity rather than aerosol sized particles which remain in the air(44). Dispersion of GFP-*E. coli* from sinks does not appear to be associated with the production of bioaerosols, that is, particles smaller than 5 μm (35, 40, 45, 46). Studies that measured air sampling lacked the resolution between aerosols and droplets(35, 40, 41). Air was sampled significantly closer (~4 inch) to the sink drain or impact point of faucet water on the sink and therefore, might have picked droplets rather than aerosols.

A consistent result from this work which is worth reemphasizing is the finding that for dispersion to occur the presence of bacteria on drain and/or bowl surface is necessary(34). When GFP-*E. coli* was inoculated into a new P-trap, dispersion was not detected using settle plates or air sampling methods. This underscores the fact that as long as the sink drain and bowl remain free of the target organisms (e.g., CPE or other antibiotic resistant Gammaproteobacteria), dispersion can be controlled. However, under favorable conditions bacteria can grow or mobilize from the P-trap into the drain piping (tailpiece) and colonize the sink drain surfaces, with the potential for a dispersion event to occur. This further underlines the importance of sanitary hygiene practices, strategic surveillance paired with hand washing only use of hand-washing sinks in the patient care environment to reduce the risk of hand-washing sink contamination by the multi-drug-resistant microorganisms that can colonize ICU patients(8). This also emphasizes the necessity to implement stricter measures to prohibit disposal of nutrients, body fluids and anything into the sinks that could be a nutrient source for maintenance of microorganism biofilms in drains(23). Dispersion from a contaminated sink reservoir can result in transmission to patients either directly or indirectly mediated through numerous contact surfaces. Herruzo and colleagues demonstrated the potential for microbial transfer from contaminated hands, which continued to disperse microorganisms after more than 10 successive contacts with surfaces (25).

The droplet dispersion load observed on settle plates was similar and consistent with our previous work(34). Total dispersion measured in corresponding experiments in the previous study was higher, which may be attributed to one or more of the following factors: i) fewer settle plates were used in the present study (22 vs 90), ii) a higher water flow rate was used in the present study (8 vs 1.8–3.0 L/min) and iii) air sampling methods performed in conjunction with the settle plate method may have captured a portion of the dispersed droplets. Settle plates were found to be a reliable method to assess the large-droplet dispersion from sinks. In this study 22 settle plates (=11.24 m^2^) were used which accounted for a defined surface area and locations on the sink counter. Dispersion could have been higher in locations of the sink counter other than those chosen in the present study, and the dispersion load recorded in this study may not be the absolute value. Of the three methods investigated for air sampling, impaction and filtration were found to be reliable and consistent. In the same amount of air sampled using impaction and filtration, comparable counts were recorded; however, air sampled using the impinger method was unable to capture the dispersion of GFP-*E. coli* under similar testing conditions.

Mannequin hands functioned as obstruction to direct impact of faucet water on the sink drain, and therefore no dispersion was detected. This rationale behind testing mannequin hands was to simulate hand washing, but in reality the water would be flowing before, after and during a hand washing event. In other words, an actual handwashing event is more dynamic than static mannequin hands and there is likely direct impact of water on the sink drain at least for brief periods when the water is running. There is also the scenario where the sinks and faucets may be used outside of hand washing (e.g. dumping liquid wastes)(5, 8, 23). This finding we think further defines and supports another important dynamic that may minimize dispersion in healthcare settings (i.e., avoid faucet water flow directly onto drains to minimize dispersion). All of these findings must be taken in the context of an experimental water stream which directly hits the drain which is outside FGI guidance but thought to be frequently found in health care sink design.

This study has several limitations. First, the dispersion experiments were not performed in a controlled environment. Each dispersion experiment lasted at least 12h, therefore it was not possible to maintain precisely the same conditions with regards to air flow velocity, air temperature, relative humidity, and bacterial and/or fungal burden in the laboratory space harboring the sinks. These parameters may have direct or indirect influence on the dispersion pattern and load recorded across experiments(47). To address this issue, we monitored the heterotrophic plate counts, relative humidity and particle concentration in the air. Particle counts recorded in the absence of faucet event (control) were higher or equal to that in the presence of faucet event (test). This observation implies that particle concentrations in the air were driven by relative humidity and/or temperature of the air. This trend was observed in all the experimental methods (Drain, P-trap inoculation and Drain colonization) (Supplemental Figure S1). In other words, particle counts were largely consistent across the day for a given experiment (control preceding test). Further particle counter used in the study could not resolve or measure particles >5μm, which defined droplet particles. Another limitation was that air samples were collected at only one location relative to the sink bowl, so it is not possible for this data set to define a “splash zone” pattern without additional measurements collected from various positions and distance from the source of dispersed organisms.

We have provided data to support the position that microorganisms will disperse from contaminated sink bowl and drain surfaces primarily as large droplets that are generated during faucet usage. These droplet-associated organisms remain viable with the potential to contaminate surfaces surrounding the sink bowl. However, it does not appear that dispersion results in the production of bioaerosols with sustained dispersion characteristics into the patient’s room space with this simulated sink design and study.

## ACKNOWLEDGEMENTS

This research was performed under a Research Collaboration Agreement between University of Virginia and the CDC. A. Mathers was supported in part by an IPA agreement between the CDC and the University of Virginia School of Medicine (15IPA1508992). The findings and conclusions in this presentation are those of the author(s) and do not necessarily represent the views of the Centers for Disease Control and Prevention. Thank you to Alexander Kallen for helpful discussions, James Matheson, Maria Burgos-Garay, and Amanda Lyons for assistance with sample processing and Will Guilford for design of the sink laboratory and automation.

## FIGURE LEGENDS

Supplemental Figure S1: Particle Counts (measured with MetOne GT-526) across Control (without faucet) and Test (with faucet) experiments of D-Drain inoculation, D7-Drainline colonization and P-P-trap inoculation methods.

## REFERENCES

1. Brooke J. 2008. Pathogenic bacteria in sink exit drains. Journal of Hospital Infection 70:198–199.

2. Hota S, Hirji Z, Stockton K, Lemieux C, Dedier H, Wolfaardt G, Gardam M. 2009. Outbreak of Multidrug-Resistant Pseudomonas aeruginosa Colonization and Infection Secondary to Imperfect Intensive Care Unit Room Design. Infect Control Hosp Epidemiol 30:25–33.

3. La Forgia C, Franke J, Hacek D, Thomson R, Robicsek A, Peterson L. 2010. Management of a multidrug-resistant Acinetobacter baumannii outbreak in an intensive care unit using novel environmental disinfection: A 38-month report. American Journal of Infection Control 38:259–263.

4. Breathnach AS, Cubbon MD, Karunaharan RN, Pope CF, Planche TD. 2012. Multidrug resistant Pseudomonas aeruginosa outbreaks in two hospitals: association with contaminated hospital waste-water systems. J Hosp Infect 82:19–24.

5. Lowe C, Willey B, O’Shaughnessy A, Lee W, Lum M, Pike K, Larocque C, Dedier H, Dales L, Moore C, McGeer A. 2012. Outbreak of extended-spectrum beta-lactamase-producing Klebsiella oxytoca infections associated with contaminated handwashing sinks(1). Emerg Infect Dis 18:1242–7.

6. Starlander G, Melhus A. 2012. Minor outbreak of extended-spectrum beta-lactamase-producing Klebsiella pneumoniae in an intensive care unit due to a contaminated sink. J Hosp Infect 82:122–4.

7. Kotsanas D, Wijesooriya WR, Korman TM, Gillespie EE, Wright L, Snook K, Williams N, Bell JM, Li HY, Stuart RL. 2013. “Down the drain”: carbapenem-resistant bacteria in intensive care unit patients and handwashing sinks. Med J Aust 198:267–9.

8. Roux D, Aubier B, Cochard H, Quentin R, van der Mee-Marquet N. 2013. Contaminated sinks in intensive care units: an underestimated source of extended-spectrum beta-lactamase-producing Enterobacteriaceae in the patient environment. J Hosp Infect 85:106–11.

9. Tofteland S, Naseer U, Lislevand J, Sundsfjord A, Samuelsen O. 2013. A Long-Term Low-Frequency Hospital Outbreak of KPC-Producing Klebsiella pneumoniae Involving Intergenus Plasmid Diffusion and a Persisting Environmental Reservoir. Plos One 8.

10. Vergara-Lopez S, Dominguez M, Conejo M, Pascual A, Rodriguez-Bano J. 2013. Wastewater drainage system as an occult reservoir in a protracted clonal outbreak due to metallo-beta-lactamase-producing Klebsiella oxytoca. Clin Microbiol Infect 19:E490–E498.

11. Knoester M, de Boer M, Maarleveld J, Claas E, Bernards A, Jonge E, van Dissel J, Veldkamp K. 2014. An integrated approach to control a prolonged outbreak of multidrug resistant Pseudomonas aeruginosa in an intensive care unit. Clin Microbiol Infect 20:O207–O215.

12. Wolf I, Bergervoet P, Sebens F, van den Oever H, Savelkoul P, van der Zwet W. 2014. The sink as a correctable source of extended-spectrum beta-lactamase contamination for patients in the intensive care unit. J Hosp Infect 87:126–130.

13. Leitner E, Zarfel G, Luxner J, Herzog K, Pekard-Amenitsch S, Hoenigl M, Valentin T, Feierl G, Grisold AJ, Hogenauer C, Sill H, Krause R, Zollner-Schwetz I. 2015. Contaminated handwashing sinks as the source of a clonal outbreak of KPC-2-producing Klebsiella oxytoca on a hematology ward. Antimicrob Agents Chemother 59:714–6.

14. Ambrogi V, Cavalie L, Mantion B, Ghiglia MJ, Cointault O, Dubois D, Prere MF, Levitzki N, Kamar N, Malavaud S. 2016. Transmission of metallo-beta-lactamase-producing Pseudomonas aeruginosa in a nephrology-transplant intensive care unit with potential link to the environment. J Hosp Infect 92:27–29.

15. Chapuis A, Amoureux L, Bador J, Gavalas A, Siebor E, Chretien ML, Caillot D, Janin M, de Curraize C, Neuwirth C. 2016. Outbreak of Extended-Spectrum Beta-Lactamase Producing Enterobacter cloacae with High MICs of Quaternary Ammonium Compounds in a Hematology Ward Associated with Contaminated Sinks. Front Microbiol 7:1070.

16. Clarivet B, Grau D, Jumas-Bilak E, Jean-Pierre H, Pantel A, Parer S, Lotthe A. 2016. Persisting transmission of carbapenemase-producing Klebsiella pneumoniae due to an environmental reservoir in a university hospital, France, 2012 to 2014. Euro Surveill 21.

17. Stjarne Aspelund A, Sjostrom K, Olsson Liljequist B, Morgelin M, Melander E, Pahlman LI. 2016. Acetic acid as a decontamination method for sink drains in a nosocomial outbreak of metallo-beta-lactamase-producing Pseudomonas aeruginosa. J Hosp Infect 94:13–20.

18. Swan JS, Deasy EC, Boyle MA, Russell RJ, O’Donnell MJ, Coleman DC. 2016. Elimination of biofilm and microbial contamination reservoirs in hospital washbasin U-bends by automated cleaning and disinfection with electrochemically activated solutions. Journal of Hospital Infection 94:169–174.

19. Zhou Z, Hu B, Gao X, Bao R, Chen M, Li H. 2016. Sources of sporadic Pseudomonas aeruginosa colonizations/infections in surgical ICUs: Association with contaminated sink trap. J Infect Chemother 22:450–5.

20. Amoureux L, Riedweg K, Chapuis A, Bador J, Siebor E, Pechinot A, Chretien M, de Curraize C, Neuwirth C. 2017. Nosocomial Infections with IMP-19-Producing Pseudomonas aeruginosa Linked to Contaminated Sinks, France. Emerging Infectious Diseases 23:304–307.

21. Baranovsky S, Jumas-Bilak E, Lotthe A, Marchandin H, Parer S, Hicheri Y, Romano-Bertrand S. 2017. Tracking the spread routes of opportunistic premise plumbing pathogens in a haematology unit with water points-of-use protected by antimicrobial filters. J Hosp Infect.

22. Bousquet A, van der Mee-Marquet N, Dubost C, Bigaillon C, Larréché S, Bugier S, Surcouf C, Mérat S, Blanchard H, Mérens A. 2017. Outbreak of CTX-M-15-producing Enterobacter cloacae associated with therapeutic beds and syphons in an intensive care unit. American Journal of Infection Control.

23. De Geyter D, Blommaert L, Verbraeken N, Sevenois M, Huyghens L, Martini H, Covens L, Pierard D, Wybo I. 2017. The sink as a potential source of transmission of carbapenemase-producing Enterobacteriaceae in the intensive care unit. Antimicrobial Resistance and Infection Control 6.

24. Lalancette C, Charron D, Laferriere C, Dolce P, Deziel E, Prevost M, Bedard E. 2017. Hospital Drains as Reservoirs of Pseudomonas aeruginosa: Multiple-Locus Variable Number of Tandem Repeats Analysis Genotypes Recovered from Faucets, Sink Surfaces and Patients. Pathogens 6.

25. Herruzo R, Ruiz G, Vizcaino MJ, Rivas L, Perez-Blanco V, Sanchez M. 2017. Microbial competition in environmental nosocomial reservoirs and diffusion capacity of OXA48 Klebsiella pneumoniae: potential impact on patients and possible control methods. J Prev Med Hyg 58:E34–E41.

26. Varin A, Valot B, Cholley P, Morel C, Thouverez M, Hocquet D, Bertrand X. 2017. High prevalence and moderate diversity of Pseudomonas aeruginosa in the U-bends of high risk units in hospital. International Journal of Hygiene and Environmental Health 220:880–885.

27. Deasy EC, Moloney EM, Boyle MA, Swan JS, Geoghegan DA, Brennan GI, Fleming TE, O’Donnell MJ, Coleman DC. 2018. Minimizing microbial contamination risk simultaneously from multiple hospital washbasins by automated cleaning and disinfection of U-bends with electrochemically activated solutions. Journal of Hospital Infection.

28. Dewi R, Tarini A, Sukrama D. 2013. Phenotypic detection of carbapenemase producing gram-negative bacteria in hospital environment abstr ASID Gram Negative Superbugs Meeting Queensland, Australia,

29. Loveday H, Wilson J, Kerr K, Pitchers R, Walker J, Browne J. 2014. Association between healthcare water systems and Pseudomonas aeruginosa infections: a rapid systematic review. Journal of Hospital Infection 86:7–15.

30. Bloomfield S, Exner M, Flemming HC, Goroncy-Bermes P, Hartemann P, Heeg P, Ilschner C, Krämer I, Merkens W, Oltmanns P, Rotter M, Rutala WA, Sonntag HG, Trautmann M. 2015. Lesser-known or hidden reservoirs of infection and implications for adequate prevention strategies: Where to look and what to look for. GMS Hyg Infect Control 10:Doc04.

31. Guyot A, Turton JF, Garner D. 2013. Outbreak of Stenotrophomonas maltophilia on an intensive care unit. J Hosp Infect 85:303–7.

32. Palmore TN, Henderson DK. 2013. Managing transmission of carbapenem-resistant enterobacteriaceae in healthcare settings: a view from the trenches. Clin Infect Dis 57:1593–9.

33. Gordon A, Mathers A, Cheong E, Gottlieb T, Kotay S, Walker A, Peto T, Crook D, Stoesser N. 2017. The Hospital Water Environment as a Reservoir for Carbapenem Resistant Organisms Causing Hospital-Acquired Infections-A Systematic Review of the Literature. Clinical Infectious Diseases 64:1435–1444.

34. Kotay S, Chai W, Guilford W, Barry K, Mathers AJ. 2017. Spread from the Sink to the Patient: in situ Study Using Green Fluorescent Protein (GFP) Expressing-Escherichia coli to Model Bacterial Dispersion from Hand Washing Sink Trap Reservoirs. Appl Environ Microbiol.

35. Schneider H, Geginat G, Hogardt M, Kramer A, Durken M, Schroten H, Tenenbaum T. 2012. Pseudomonas aeruginosa outbreak in a pediatric oncology care unit caused by an errant water jet into contaminated siphons. Pediatr Infect Dis J 31:648–50.

36. Fusch C, Pogorzelski D, Main C, Meyer C, el Helou S, Mertz D. 2015. Self-disinfecting sink drains reduce the Pseudomonas aeruginosa bioburden in a neonatal intensive care unit. Acta Paediatrica 104:e344–e349.

37. Pitten FA, Panzig B, Schröder G, Tietze K, Kramer A. 2001. Transmission of a multiresistant Pseudomonas aeruginosa strain at a German University Hospital. Journal of Hospital Infection 47:125–130.

38. Siegel JD, Rhinehart E, Jackson M, Chiarello L. 2007. 2007 Guideline for Isolation Precautions: Preventing Transmission of Infectious Agents in Health Care Settings. Am J Infect Control 35:S65–164.

39. WHO. 2014. Infection prevention and control of epidemic-and pandemic-prone acute respiratory infections in health care: WHO Guidelines. World Health Organization, Geneva. http://apps.who.int/iris/bitstream/10665/112656/1/9789241507134eng.pdf?ua¼1.

40. Kramer A, Daeschlein G, Niesytto C, Sissoko B, Sütterlin R, Blaschke M, Fusch C. 2005. Contamination of sinks and emission of nosocomial gram negative pathogens in a NICU-outing of a reservoir as risk factor for nosocomial colonization and infection Umweltmed Forsch Prax 10.

41. Doring G, Horz M, Ortelt J, Grupp H, Wolz C. 1993. Molecular epidemiology of Pseudomonas aeruginosa in an intensive care unit. Epidemiol Infect 110:427–36.

42. Zemouri C, de Soet H, Crielaard W, Laheij A. 2017. A scoping review on bio-aerosols in healthcare and the dental environment. Plos One 12.

43. Haig C, Mackay W, Walker J, Williams C. 2016. Bioaerosol sampling: sampling mechanisms, bioefficiency and field studies. Journal of Hospital Infection 93:242–255.

44. Morawska L. 2006. Droplet fate in indoor environments, or can we prevent the spread of infection? Indoor Air 16:335–347.

45. Eames I, Shoaib D, Klettner C, Taban V. 2009. Movement of airborne contaminants in a hospital isolation room. Journal of the Royal Society Interface 6:S757–S766.

46. Stetzenbach L. 2002. Introduction to aerobiology. In Hurst C, Crawford R, Knudsen G, McInerney M, Stetzenbach L (ed), Manual of environmental microbiology, 2nd ed. ASM Press, Washington, DC.

47. Mohr A. 2002. Fate and transport of microorganisms in the air, p 827–838. In Hurst C, Crawford R, Knudsen G, McInerney M, Stetzenbach L (ed), Manual of Environmental Microbiology, 2nd ed. ASM Press, Washington, DC.

